# Nearly unbiased estimator of contemporary effective population size using within-cohort sibling pairs incorporating parental and non-parental reproductive variations

**DOI:** 10.1101/631085

**Authors:** Tetsuya Akita

**Author notes:** Corresponding author:* National Research Institute of Fisheries Science, Japan Fisheries Research and Education Agency, 2-12-4 Fukuura, Kanazawa, Yokohama, Kanagawa, 236-8648, Japan.

## Abstract

In this study, we developed a nearly unbiased estimator of contemporary effective mother size in a population, which is based on a known maternal half-sibling relationship found within the same cohort. Our method allows for variance of the average number of offspring per mother (i.e., parental variation, such as age-specific fecundity) and variance of the number of offspring among mothers with identical reproductive potential (i.e., non-parental variation, such as family-correlated survivorship). We also developed estimators of the variance and coefficient variation of contemporary effective mother size and qualitatively evaluated the performance of the estimators by running an individual-based model. Our results provide guidance for (i) a sample size to ensure the required accuracy and precision when the order of effective mother size is available and (ii) a degree of uncertainty regarding the estimated effective mother size when information about the size is unavailable. To the best of our knowledge, this is the first report to demonstrate the derivation of a nearly unbiased estimator of effective population size; however, its current application is limited to effective mother size and situations in which the sample size is not particularly small and maternal half-sibling relationships can be detected without error. The results of this study demonstrate the usefulness of a sibship assignment method for estimating effective population size; in addition, they have the potential to greatly widen the scope of genetic monitoring.

## Introduction

Contemporary effective population size, which is sensitive to ecological time-scale events, has become recognized as an informative parameter in a focus population, especially in the context of conservation biology and wildlife management (Luikart *et al*., 2010). There are several methods for estimating contemporary effective population size from genetic markers, such as the temporal method (Nei and Tajima, 1981), heterozygote excess method (Pudovkin *et al*., 1996), molecular coancestry method (Nomura, 2008), linkage-disequilibrium method (Waples, 2006), and kinship assignment method (Wang, 2009). At present, it is known that values estimated by these methods display large uncertainties and/or biases under conditions such as small sample size, small marker numbers, and large effective population size; thus, a widely applicable method is required (Wang *et al*., 2016).

Owing to rapid developments in genotyping technology, a large number of genetic markers, including thousands of genome-wide single nucleotide polymorphisms, have become available for analyzing population structure and demography. As a result, a more accurate estimation of contemporary effective population size can be obtained by, for example, more accurately assigned kinships (Wang *et al*., 2016). In addition, the recently developed theory of estimation of absolute adult number, which is based on sampled kinship pairs and known as the close-kin mark-recapture (CKMR) method (Bravington *et al*., 2016a,b; Skaug, 2017; Hillary *et al*., 2018), makes it possible to use a full-sibling (FS) or half-sibling (HS) pair; this involves many more DNA markers for detection than a parent–offspring pair. It should be noted that the CKMR method is designed to minimize the effect of reproductive variance originating from unmodeled covariates, such as avoiding the use of sibling pairs sampled from the same cohorts; meanwhile, reproductive variance strongly affects the estimation of contemporary effective population size.

Reproductive variance has two components. The first component is variation in age, size, and other factors, which affects average fecundity and originates from differences in life-history parameters (Felsenstein, 1971). For example, in the case of teleost species that have a long life span, the number of eggs produced by a mother (i.e., annual fecundity) is determined by her body size; thus, there is considerable variation in reproduction among mothers. The second component is variation in reproduction among parents of the same age or size. An extreme case reflecting this variation is referred to as the “Sweepstakes Reproductive Success (Hedgecock and Pudovkin, 2011),” in which only several families reproduce successfully. This phenomenon has received much attention not only for elucidating the ecology of species that display highly variable early life mortality (i.e., type-III life history) but also for providing an opportunity to test the applicability of the multiple-merger coalescent model, a recently developed theory in population genetics (Tellier and Lemaire, 2014; Eldon *et al*., 2016). Addressing the two aforementioned types of variance together can provide insights for interpreting estimated values of effective population size.

In this paper, we propose a new method for estimating the contemporary effective mother size in a population. This approach is based on the number of maternal HS (MHS) pairs found within the same cohort and on modeling that explicitly incorporates overdispersed reproduction, assuming that kinships are genetically detected without any error. Our model divides reproductive variance into two types of variations: (i) age- or size-specific differences in mean fecundity (referred to as “parental variation,”) and (ii) unequal contributions by mothers of the same age or size to the number of offspring at sampling (referred to as “non-parental variation.”) First, we formulate the distribution of offspring number under the two types of variations. Second, we analytically derive the probability that two randomly selected individuals found in the same cohort share an MHS relationship. Third, we determine a nearly unbiased estimator of contemporary effective mother size and its relative estimators. Finally, we investigate the performance of the estimators by running an individual-based model. Our modeling framework may be applied to diverse animal species; however, the description of the model focuses on fish species, which are currently the best candidate target of our proposed method.

## Theory

All symbols used in this paper are summarized in Table 1.

**Table 1:** List of mathematical symbols in main text

|  |  |
| --- | --- |
| $n$ | Sample number of offspring |
| $n_{\text{pair}}$ | Number of pairs in a sample ($= {}_n\text{C}_2$) |
| $N$ | Number of mothers in the population when sampled offspring are born |
| $N_e$ | Effective number of mothers in the population |
| $\phi$ | Overdispersion parameter under negative binomial reproduction |
| $\lambda_i$ | Expected number of surviving offspring of mother $i$ at sampling |
| $f(\lambda)$ | Frequency of $\lambda$ for all mothers. |
| $k_i$ | Number of surviving offspring born to mother $i$ |
| $H$ | Number of maternal–half-sibling pairs found in samples |
| $\pi$ | Probability that a randomly selected pair (two offspring) share a maternal half-sibling relationship |
| $c$ | Combined effect of deviation from the Poisson<br>($= (1 + \phi^{-1})\mathbb{E}[\lambda^2]/\mathbb{E}[\lambda]^2$) |
| $\hat{N}_{e,0}$ | Moment estimator of $N_e$ |
| $\hat{N}_{e,1}$ | Nearly unbiased estimator of $N_e$ |
| $\hat{v}$ | Estimator of $\mathbb{V}[\hat{N}_{e,1}]$ |
| $\hat{c}v$ | Estimator of $\mathbb{CV}[\hat{N}_{e,1}]$ |
| $b_{\text{mean}}$ | Bias of $\hat{N}_{e,1}$ |
| $b_{\text{var}}$ | Bias of $\hat{v}$ |

### Hypothetical population and sampling scheme

Here, we suppose that there is a hypothetical population consisting of *N* mothers and that there is no population subdivision or spatial structure. In this paper, a mature female is referred to as a mother even if she does not produce offspring. For the detection of MHS pairs, *n* offspring within the same cohort are simultaneously and randomly sampled in the population. For mathematical tractability, we assume that there is only one spawning ground in which the mothers remain for the entire spawning season.

In our modeling framework, if an MHS pair also shares a paternal HS (PHS) relationship, the pair is considered to be an MHS pair (i.e., the FS relationship is assigned as MHS relationship). In addition, we assume that there are sufficient opportunities for every mother to mate; therefore, the behavior of fathers does not affect the reproductive output of mothers. These simplifications can avoid the complex interactions between mothers and fathers in one breeding season. The limitations of these simplifications, as well as the technical difficulties of distinguishing an MHS pair from a PHS pair, are addressed in the **Discussion** section.

### Reproductive potential and its variation (parental variation)

Here, we introduce the concept of the reproductive potential of mother *i* (*i* = 1,2, …,*N*), which is defined as the expected number of surviving offspring at sampling time, denoted by *λ_i_*. The reproductive potential is determined by several factors, including the mother’s age, weight, or residence time on the spawning ground, and it is allowed to vary across mothers. In this study, this variation is referred to as parental variation. It should be noted that the magnitude of this parameter (*λ_i_*) includes information about the survival rate of the offspring, the number of days after egg hatching, and the egg number; this implies that the parameter reflects the sample timing.

### Non-parental variation

In addition to parental variation, the variation in reproduction among mothers with the same reproductive potential, referred to here as non-parental variation, is also incorporated into the model, resulting in a large variation in the fertility of the mothers. As the magnitude of the variance increases, the number of successful mothers producing offspring that avoid early life mortality decreases, leading to a situation in which offspring derived from the same mother have highly correlated early life survival probabilities. This situation requires careful consideration of the probability that two offspring share an MHS relationship. **Fig. 1** presents a schematic representation of the effects of such family-correlated survival on kinship relationships in a population, which are exemplified in iteroparous teleost species. Older mother are more likely to produce a larger number of offspring, as annual fecundity (i.e., number of eggs, represented by a gray circle) increases with age. However, due to family-correlated survivorship after eggs hatching, the probability that two offspring (i.e., at the larva or juvenile stage, represented by a closed circle) have an MHS relationship is higher (e.g., 53 MHS pairs in **Fig. 1b**) than in a situation with independent survival (e.g., 32 MHS pairs in **Fig. 1a**). In other words, MHS pairs have significantly higher or lower collective chances for survival. Non-parental variation may occasionally overshadow the effect of parental variation; however, the average number of offspring per mother is higher for an older mother if the probability of being a successful mother is not biased among mothers.

**Figure 1:**
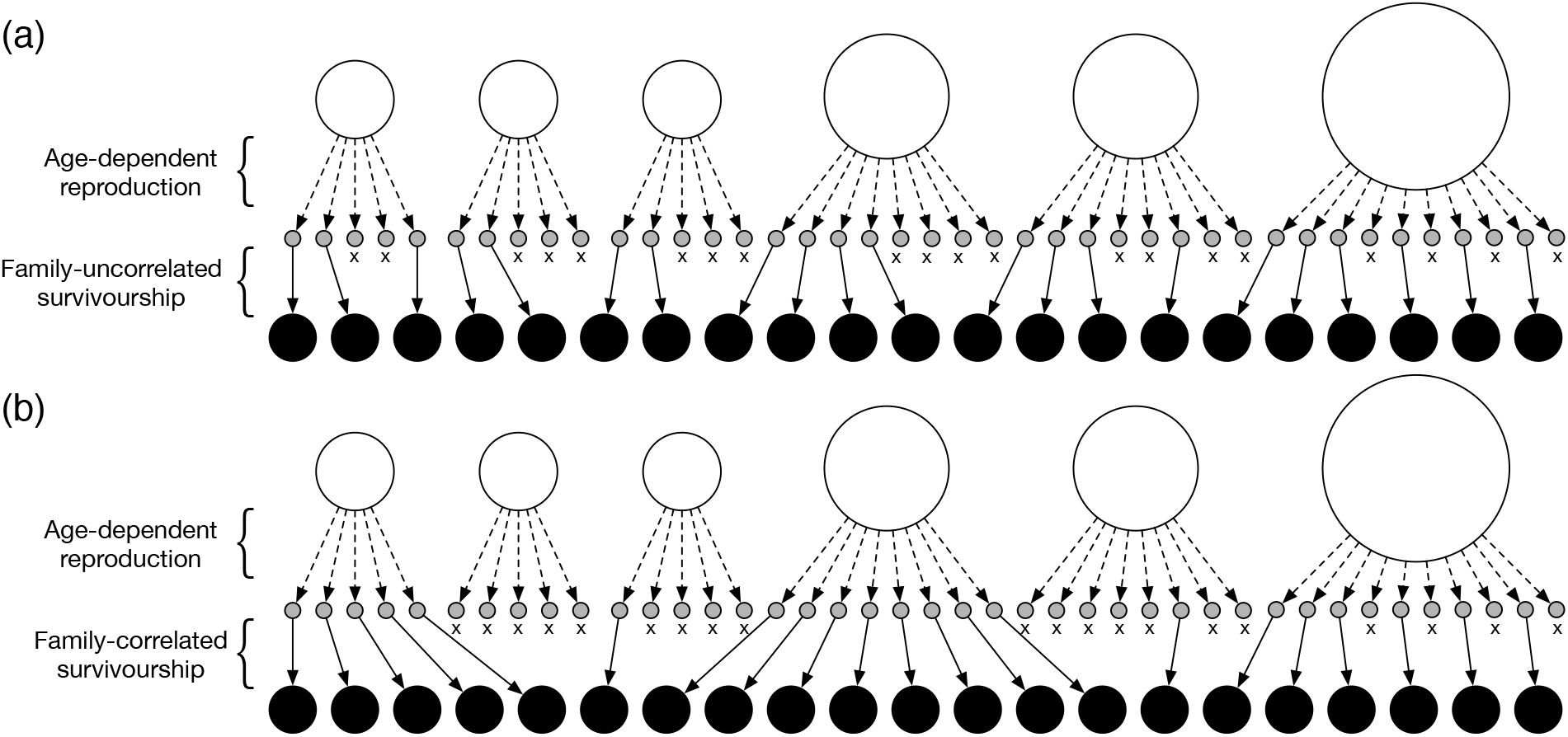
Example of relationships between mothers and their offspring number for only parental variation (a) and both paretal- and non-parental variation (b). *N* = 6 and 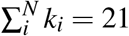. Open, gray, and black circles represent mothers, their eggs, and their offspring, respectively. The area of an open circle indicates the degree of reproductive potential of each mother (i.e., *λ_i_*). Dotted and thin arrows show mother–egg and egg–offspring relationships, respectively. The x symbol indicates a failure to survive at sampling.

### Distribution of offspring number

In attempting to incorporate both parental and non-parental variation, it is useful to employ a highly skewed distribution of offspring number. In this study, we use a negative binomial distribution, which is applicable to deviation from the Poisson variance (i.e., overdispersed offspring number with a variance greater than the mean). In the overdispersed reproduction, whether the *l*th offspring is born to the ith mother depends on the probability that the *m*th offspring is the ith mother and is thus the *l*th offspring with an MHS relationship.

Let *k_i_* be the number of surviving offspring of mother *i* at sampling. Given the expected number of offspring *λ_i_, k_i_* is assumed to follow a negative binomial distribution by a conventional parametrization,

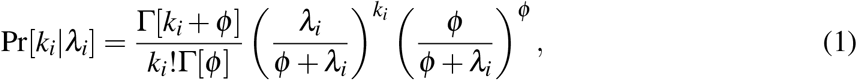

where *ϕ* (> 0) is the overdispersion parameter describing the degree of non-parental variation (Akita, 2018). At present, *ϕ* is assumed to be constant across mothers, whereas the expected number of surviving offspring (*λ_i_*) is variable across mothers. The mean and variance of this distribution are *λ_i_* and 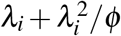, respectively. In the limit of infinite *ϕ*, this distribution becomes a Poisson distribution as follows:

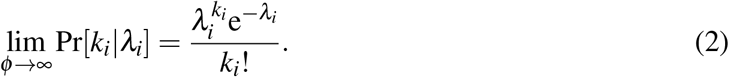

We assume that *λ_i_* is independent and identically distributed with a density function *f* (*λ*), which produces parental variation. The shape of the density function is often complex but may be described by information such as the mother’s weight composition in the population. The specific form of *f* (*λ*) is provided in **Appendix A** and is used for verifying the theory developed in this paper. As explained in the next subsection, the theory does not require this specific form; it only requires the ratio of the second moment to the squared first moment (i.e., 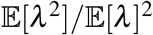).

For illustrative purposes, we demonstrate how parental and non-parental variation skew the offspring distribution. **Figure 2a** presents a histogram of *f* (*λ*) for two cases–one in which parental variation is scarcely observed (represented by a black bar) and the other in which it is moderately observed (represented by a gray bar). The cases can be controlled by changing the parameters affecting the shape of *f* (*λ*), such as the allometric growth parameter (defined by *β*; for details, see **Appendix A**). If both parental and non-parental variations are very small, *k* has as a Poisson distribution (dotted line in **Fig. 2b**), as noted above. When there is no parental variation, non-parental variation skews the distribution of *k* (thin and bold lines in **Fig. 2b**), and vice versa (dotted line in **Fig. 2c**).

**Figure 2:**
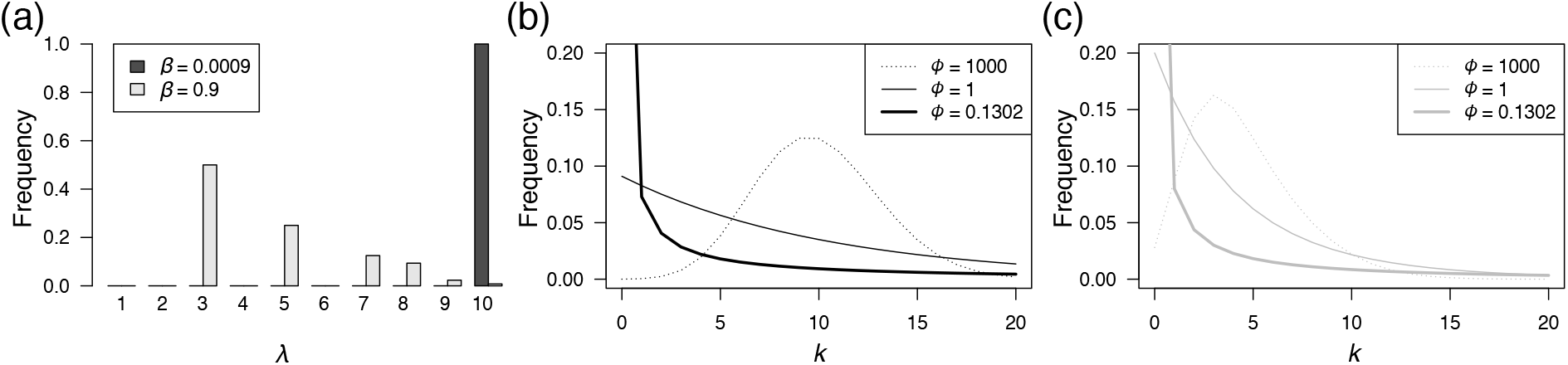
(a) Histogram of *f* (*λ*) assuming fish species with a relatively low *β* (denoted by black bar, 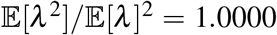) and high *β* (denoted by gray bar, 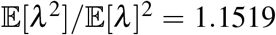). (b), (c) Marginal distribution of *k* for several values of *ϕ* (see legend). (b) *β* = 0.0009; Pr[*k* = 0] with *ϕ* = 0.1302 equals 0.57. (c) *β* = 0.9; Pr[*k* = 0] with *ϕ* = 0.1302 equals 0.63. Details of *f* (*λ*) are provided in **Appendix A**.

In this study, we selected parameter 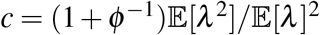 (see the next subsection) to be 1 and 10 for comparison with the results in the main text. These two values represent two extreme cases and can be derived from the parameter set (*ϕ, β*) = (1000,0.0009) (dotted line in **Fig. 2b**) and (0.1302,0.9) (bold line in **Fig. 2c**), respectively. It should be noted that in the latter case, the offspring distribution is highly skewed: 63% of mothers cannot produce surviving offspring at sampling, and 6% of mothers produce more than 20 offspring 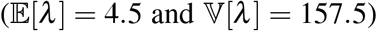. Other parameter values used in *f* (*λ*) are provided in **Appendix A**.

### MHS probability among randomly selected individuals

We have derived the probability that two offspring share an MHS relationship with an arbitrary mother as follows (see **Appendix B**):

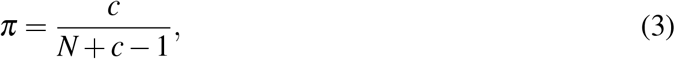

where

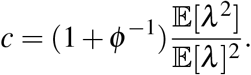

Equation 3 explicitly contains the two variations (i.e., parental variation and non-parental variation) that determine the degree of deviation from the Poisson distribution. When *λ* is constant across mothers, 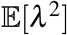 equals 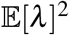 and then *π* becomes (1 + *ϕ*^−1^)/(*N* + *ϕ*^−1^), which appears in Eq. 7 in Akita (2018). In addition, as *ϕ* → ∞, (1 + *ϕ*^−1^)/(*N* + *ϕ*^−1^) converges to 1/*N*, which corresponds to the Poisson variance of *k_i_* for all mothers in a population. The effect of the two factors causing a deviation from the Poisson distribution can be combined as parameter *c* (≥ 1). Hereafter, “overdispersion” is referred to as the distribution of the number of offspring resulting from this combined effect.

When *N* is provided, *π* increases with an increase in *c*, suggesting that a randomly selected pair is more likely to share an MHS relationship under greater overdispersion. **Figures S1a–d** (**Supplementary Information**) illustrate the theoretical curve and the simulation results of *π* with *N* = 100 and 10,000 as a function of *ϕ* or 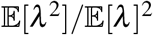. These figures demonstrate that the approximation in Eq. 3 works well for the investigated function *f* (*λ*).

### Effective mother size and census size

We have defined the effective mother size as follows:

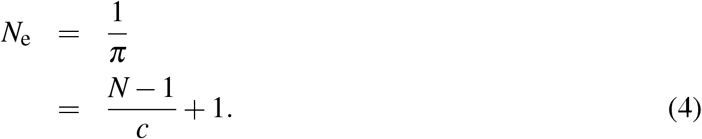

This definition is similar to the coalescent effective population size (Nordborg and Krone, 2002), as the probability of sharing an MHS relationship (*π*) is identical to the probability that two individuals coalesce into a mother in the previous breeding season. It should be noted that when sampling from a single cohort in a population with overlapping generations, the effective mother size in our definition corresponds to the effective breeding mother size, which produces a single cohort.

Using Eq. 4, the ratio of the effective mother size to census size can be written as follows:

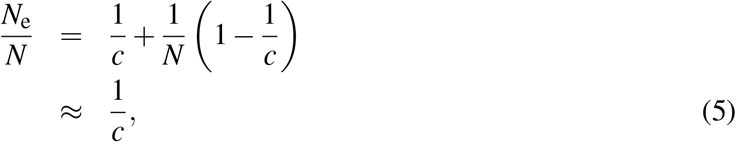

where *N* ≫ 1 is assumed for the purpose of approximation.

### Statistical properties of MHS pair number

In this subsection, given the unconditional probability that two offspring share an MHS relationship (Eq. 3), we consider the distribution of the number of MHS pairs and its statistical properties. Let *H* be the number of MHS pairs found in an offspring sample of size *n*. First, we derive the approximate distribution of *H* for a situation in which overdispersion does not exist (i.e., *c* = 1). Second, we evaluate whether the derived distribution of *H* for the non-overdispersed case is applicable to the overdispersed case (i.e., *c* > 1).

If overdispersion does not exist (i.e., *c* = 1), drawing an MHS pair from a randomly selected pair in a sample is considered a Bernoulli trial. Thus, *H* follows a hypergeometric distribution, which is a function of the sample size of the offspring, the total number of offspring in the population, and the total number of MHS pairs in the population. However, in the setting of this study, the latter two components are random variables, thus creating a complex situation for deriving the exact formulation (Akita, 2018). Therefore, assuming that the total number of MHS pairs in the population is much higher than the number of pairs in a sample _Σ*k*_C_2_ ≫ _*n*_C_2_ the distribution is approximated by a binomial form as follows:

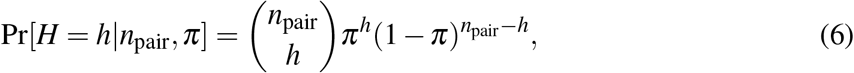

where *n*_pair_ is the number of pairs in a sample (= _*n*_C_2_). For practical purposes, the condition _Σ*k*_C_2_ ≫ _*n*_C_2_ may be acceptable. The theoretical expectation of *H* is

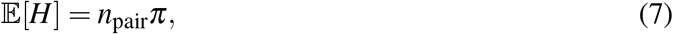

and the variance is

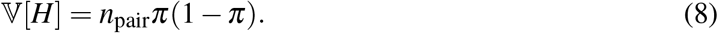

**Figures S2a and S2b** (**Supplementary Information**) illustrate the accuracy of the theoretical prediction for the expectation and the variance of *H* under the Poisson variance as a function of *n*, respectively. For the investigated parameter, the prediction is demonstrated to be highly accurate.

If overdispersion exists (i.e., *c* > 1), drawing an MHS pair is no longer a Bernoulli trial. For example, an individual that is born to a relatively successful mother has a greater probability of an MHS relationship with other individuals. Therefore, a hypergeometric/binomial form is not appropriate for the distribution of *H*. As illustrated in **Fig. S2d** (**Supplementary Information**), the binomial variance (Eq. 8) is downwardly biased from the observed variance of *H* when *n* increases. The theoretical evaluation is relatively complex and is left for future research. However, for the investigated parameter set, the expected value is well approximated by Eq. 7 (**Fig. S2c** in **Supplementary Information**), assuming independent comparisons. The rationale may be that the MHS probability in a pair, *π* (Eq. 3), includes the effect of overdispersion. Next, on the basis of an accurate approximation of 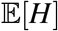 in the case of overdispersion, we provide the estimator of *N*_e_ from the observed number of MHS pairs in a sample.

### Moment estimator of *N*_e_ from observed number of MHS pairs

By removing *π* in Eqs. 4 and 7, *N*_e_ can be written as a function of *c*, *n*_pair_, and 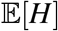. The observed number of MHS pairs in a sample is defined by *H*_obs_, and 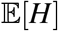 is replaced by *H*_obs_, generating the moment estimator of *N*_e_:

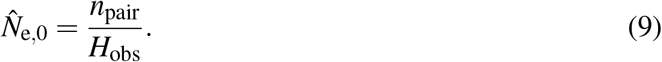

In this paper, a “hat” indicates the estimator of a variable. This relationship can be written as follows:

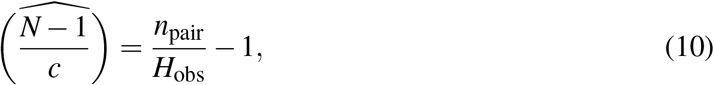

indicating that *N* and *c* cannot be estimated simultaneously from the number of observed MHS pairs.

Assuming that *H* follows a binomial distribution, the estimator corresponds to the maximum likelihood estimator of *N*_0_ (see **Appendix C**). There are two drawbacks to using this estimator. First, the value of 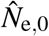 becomes inflated when no MHS pairs are observed in a sample (i.e., *H*_obs_ = 0). This leads to a situation in which an individual-based model (IBM) frequently generating zero MHS pairs is not available for statistical evaluation. Second, even if an MHS pair is detected in a sample, it is likely that 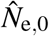 is strongly biased (see **Appendix C**). Therefore, an improved estimator is necessary for the purpose of appropriate evaluation and higher accuracy for a wide parameter range.

### Nearly unbiased estimator of *N*_e_

We have derived an alternative estimator of *N*_e_ (see **Appendix D**) as follows:

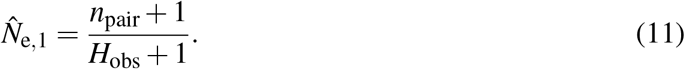

The derivation process is similar to that of the nearly unbiased estimator of adult number in a population using the mark-recapture method (Chapman, 1951). The bias of 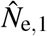 is defined by *b*_mean_, which is given by

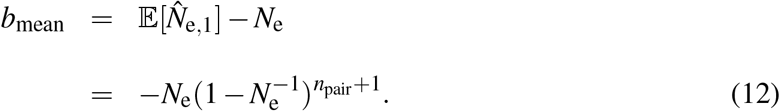

It should be noted that 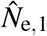 is downwardly biased; however, this bias may be ignored for a wider range of parameters than 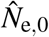 (see details in the **Results** section), which allows 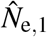 to be called a nearly unbiased estimator.

We also determined the estimator of 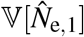, given by

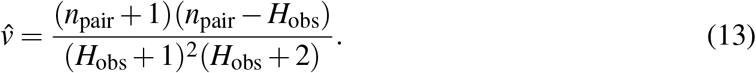

The derivation process is similar to that in Seber (1970) (see **Appendix E** for details). The bias of 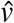 is defined by *b*_var_, which is given by

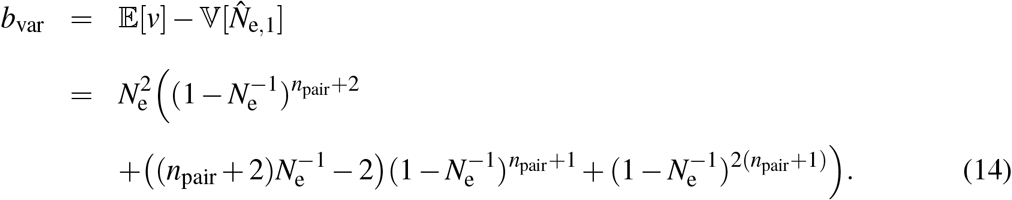

Finally, we consider the estimator of the coefficient of variation of 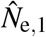. A method similar to the derivation of 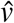 (i.e., searching for a formula such that its expectation approximates 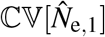 was overly complex for the estimator; instead, using Eqs. 11 and 13, we defined the estimator as follows:

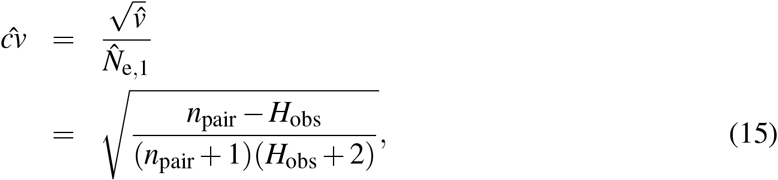

Roughly speaking, 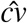 is approximated by 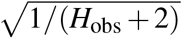 because *n*_pair_ ≫ *H*_obs_, which is similar to an approximate lower bound on the coefficient of variation of 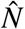, as presented in Bravington *et al*. (2016b).

### Individual-based model (IBM)

To evaluate the performance of the estimators (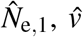, and 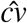), we developed an IBM that tracks kinship relationships. The population structure was assumed to be identical to that in the development of the estimators. The population consisted of mothers and their offspring and was assumed to follow a Poisson or negative binomial reproduction. The expected number of surviving offspring of a mother followed the density distribution *f* (*λ*) (see **Appendix A**). It should be noted that the overdispersion parameter (*c*) was calculated from *ϕ* and *f* (*λ*). Each offspring retained the ID of its mother, making it possible to trace an MHS relationship.

Given a parameter set (*N, c, n, ϕ*, and parameters that determine *f* (*λ*)), we simulated a population history in which *N* mothers generated offspring; this process was repeated 100 times. For each history, the sampling process was repeated 1000 times, acquiring 100,000 data points that were used to construct the distribution of 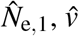, and 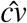 for each parameter set. *N*_e_ was calculated from *N* and *c* (Eq. 4).

## Results

We evaluated the performance of the estimators (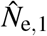, and 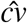) for a situation in which the number of mothers, *N*, and the combined effect of deviation from the Poisson, *c*, were unknown. The parameter values were changed for *N* (10^4^, 10^5^, and 10^6^) and *c* (1 and 10). We primarily addressed the number of samples (*n*) required to provide adequate accuracy and precision under a given parameter set (*N* and *c*).

First, we evaluated the accuracy of 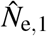 based on its bias (*b*_mean_). For a given *N*_e,1_, the absolute value of the bias is represented by a solid line in **Figs. 3a** (*N*_e,1_ = 10,000) and **3b** (*N*_e,1_ = 100,000) as a function of *n*. For comparison, the bias of 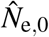 (see **Appendix C**) is represented by a dotted line. It is evident that the absolute value of the bias is smaller in 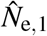 than in 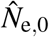. Hereafter, we use 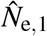 as the estimator of effective mother size.

**Figure 3:**
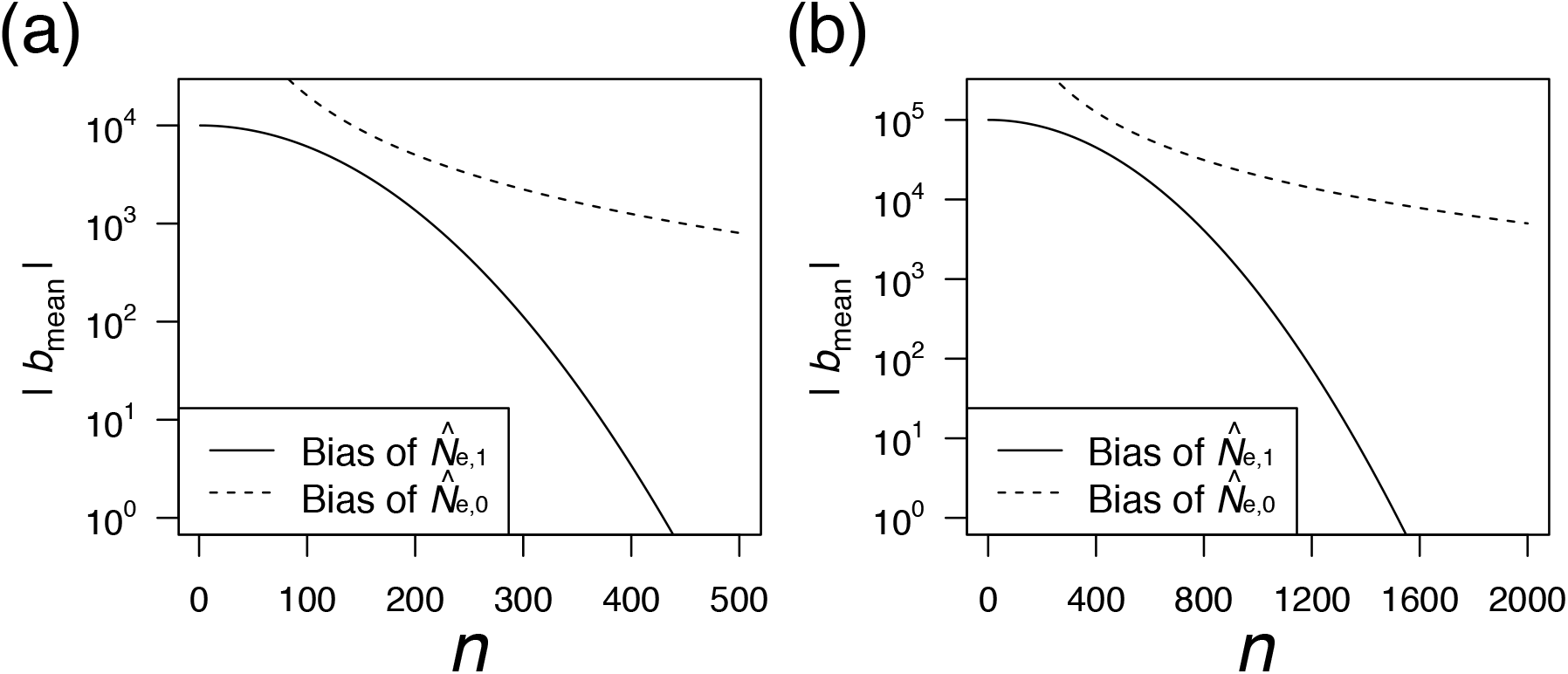
Absolute bias of 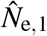 (represented by solid line) and 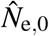 (represented by dotted line) as a function of *n*. (a) *N*_e_ = 10,000. (b) *N*_e_ = 100,000.

As expected, |*b*_mean_| decreases with *n*, as a larger *n* leads the term 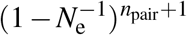 in *b*_mean_ to vanish more quickly. The requisite sample size (*n*) with a small bias of less than 1% is approximately 300 for *N*_e_ = 10,000 (|*b*_mean_| < 100; see **Fig. 3a**) and 1000 for *N*_e_ = 100,000 (|*b*_mean_| < 1000; see **Fig. 3b**). The results of the IBM support the above prediction. **Figure 4** illustrates the average value of 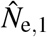 (represented by open circles) with a 95% confidence interval (CI), which is obtained from the IBM. As expected, the average value of 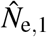 downwardly deviates from *N*_e_ for a relatively small sample size (*n*) satisfying |*b*_mean_| ≫ 1. As *n* increases, the average value of 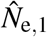 approaches a true *N*_e_ (represented by a dotted line in **Fig. 4**).

**Figure 4:**
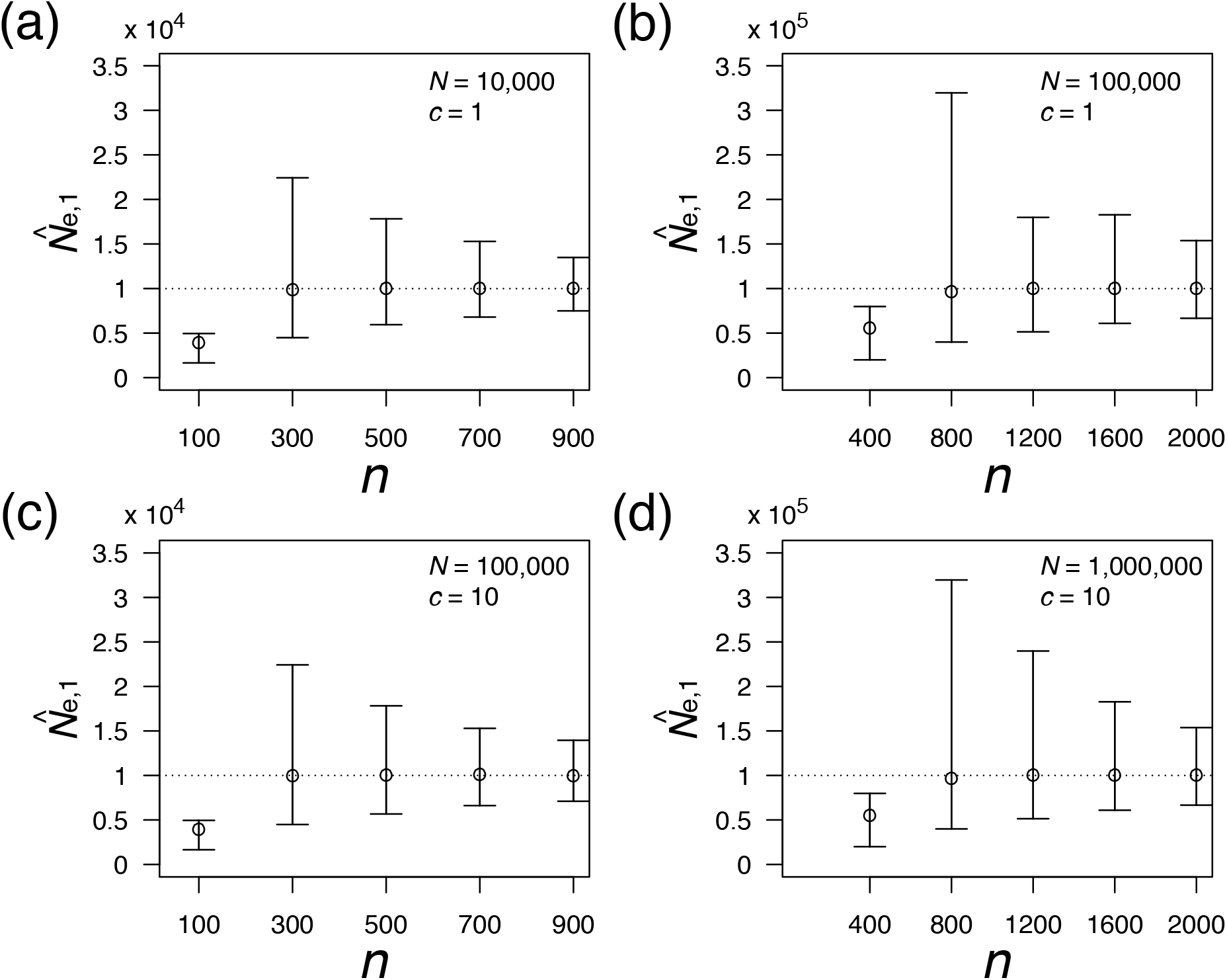
Accuracy and precision of 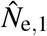 as a function of *n*. Open circles represent means with 95% CIs. A dotted line indicates the true value of *N*_e_. The value of parameters (*N* and *c*) is indicated in the legend. (a), (c) *N*_e_ ≈ 10,000. (b), (d) *N*_e_ ≈ 100,000.

Next, we evaluated the precision of 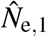. As illustrated in **Fig. 4**, the precision of 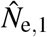 for a change in *n* behaves in a complex manner. For the investigated parameter set, we determined that the degree of precision holds under different combinations of *N* and *c* if the value of *N*_e_ is fixed (*N*_e_ ≈ *N*/*c* equals 10,000 in **Figs. 4a and 4b** and 100,000 in **Figs. 4c and 4d**); this suggests that the level of uncertainty is roughly determined by *N*_e_. Although the lower limit of the CI monotonically increases with *n*, the upper limit of the CI has a peak at the point at which the average 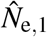 is very close to the true *N*_e_. Near this point, the range of the CI is large, and 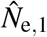 is asymmetrically distributed with a longer tail on the large side (e.g., *n* = 800 in **Figs. 4b and 4d**). As *n* increases beyond this point, the range of the CI decreases, and the shape of the distribution asymptotically becomes symmetric.

We then evaluated the accuracy of 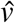. Theoretically, the bias of 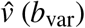 was determined to have a peak at a certain value of *n*, as illustrated in **Figs. S3a and S3b** (**Supplementary Information**). **Figures 5a and 5c** present the ratio of the average 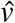 to the variance of 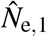 for different combinations of (*N,c*) with fixed *N*_e_, which is obtained from the IBM. If the ratio is close to 1, 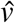 is deemed an estimator of unbiasedness. When *n* approaches zero, the ratio becomes inflated (e.g., *n* = 100 in **Fig. 5a**) although *b*_var_ also approaches zero (**Figs. S3a and S3b**). This inconsistency for a small *n* (i.e., overestimation) may result from the bias of 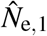. As *n* increases, the ratio approaches 1 when *c* = 1 but less than 1 when *c* > 1 (*c* = 10 in **Fig. 5a**), suggesting that the property of unbiasedness holds only under the Poisson variance; however, the degree of this bias is not very high for a relatively large *N*_e_ (*c* = 10 in **Fig. 5b**). In other words, the accuracy of 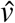 is not solely determined by the level of *N*_e_. This inconsistency (i.e., underestimation) may result from the assumption that the correlation between pairs can be ignored and thus that the number of HS pairs in the sample follows a binomial distribution (Eq. 6).

**Figure 5:**
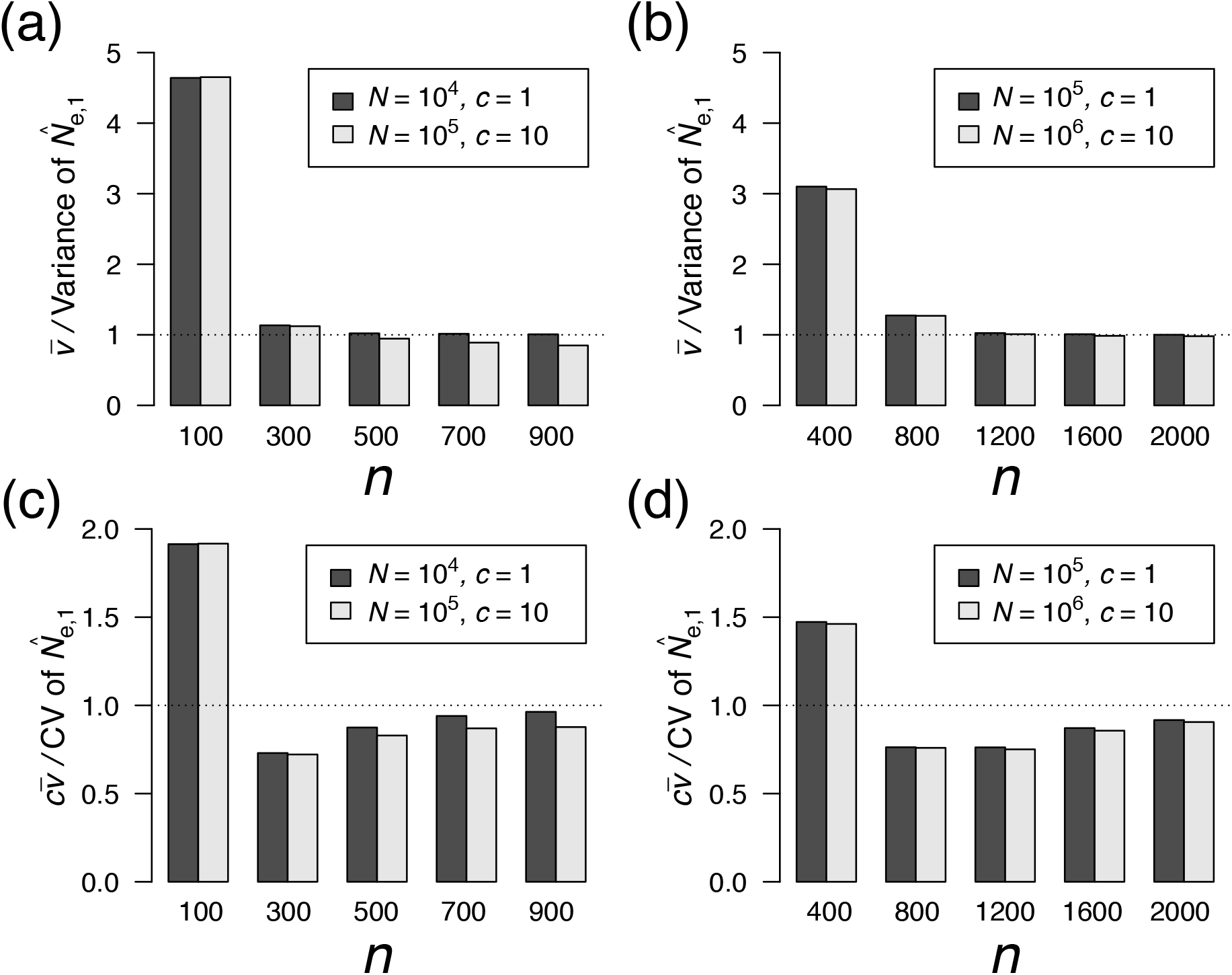
(a), (b) Ratio of the average 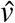 to the variance of 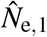 as a function of *n*. (c), (d) Ratio of the average 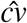 to the coefficient of variation of 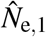 as a function of *n*. The value of parameters (*N* and *c*) is indicated in the legend. (a), (c) *N*_e_ ≈ 10,000. (b), (d) *N*_e_ ≈ 100,000.

Finally, we evaluated the accuracy of 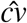. **Figures 5c and 5d** illustrate the ratio of the average 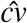 to the coefficient of variation of 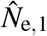, which is obtained from the IBM. As expected, the property of the estimator is similar to that of 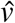, as 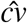 is defined by using 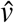 (Eq. 15). The ratio becomes inflated for small *n*; as *n* increases, the ratio approaches 1 when *c* = 1 but is less than 1 when *c* > 1 (i.e., underestimation); however, the relationship between the degree of bias and the level of *N*_e_ is unclear.

## Discussion

In this study, we theoretically developed a nearly unbiased estimator of the number of effective mothers in a population 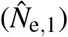, the estimator of its variance 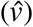, and its coefficient variation 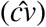, which are based on the known MHS relationships found within a single cohort. The performance of the estimators (accuracy and precision) was quantitatively evaluated by running an IBM. Our modeling framework allows for two types of reproductive variation; variance of the average offspring number per mother (parental variation) and variance of the offspring number across mothers with the same reproductive potential (non-parental variation). The former is related to age- or size-dependent reproductive potential, whereas the latter is related to family-correlated survival, both of which can result in a skewed distribution of offspring number. These two effects are summarized into one parameter (*c*) in the framework. Our estimators can be calculated from sample size (*n*) and the observed number of MHS pairs (*H*_obs_) and do not require other parameters, such as adult mother size (*N*) or the degree of overdispersed reproduction (*c*). The rationale for this is that the observed number of MHS pairs contains information about these parameters.

To estimate the number of effective mothers (*N*_e_), our theoretical results provide guidance for a sample size to ensure the required accuracy and precision, especially if the order of the number of effective mothers is approximately known. For example, when the effective number of mothers is within 10^4^–10^5^, sampling 500 offspring falls within the range of accuracy of the estimation with a 0–30% bias (Eq. 12 and **Fig. 3**). Even if there is no information about the effective number of mothers, the coefficient of variation of the estimated number can be estimated 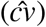 when the sample size is above a given level (**Figs. 5c and 5d**). Although the estimator of the variation of the number of mothers 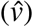 is relatively accurate for the investigated parameter set (**Figs. 5a and 5b**), the present estimator of the coefficient variation is systematically biased; thus, improvements in accuracy are left for future research. In addition, the main contribution of this study is to provide nearly unbiased estimators; theoretical compatibility and performance comparisons between our results and those of existing methods are left in future work.

Our modeling framework is presented in the context of the sibship assignment (SA) method, which defined a kinship-oriented estimation of effective population size (Wang, 2009; Waples and Waples, 2011). The original theory of the SA method was developed by Wang (2009), and it can perform the estimation of effective population size from HS and FS probabilities, which are calculated by the number of HS and FS pairs in a sample. To the best of our knowledge, in SA method, there is little theoretical foundation for a nearly unbiased estimator of effective population size and its variance; therefore, the evaluation of accuracy and precision has relied on numerical methods. In this study, we analytically obtained nearly unbiased estimators, although their application is limited to the estimation of effective mother size and the case in which MHS can be perfectly distinguished from PHS and other relatives. The latter limitation may be overcome to some extent with the use of a hypervariable region in the mitochondrial genome and/or sex-linked markers. It should be noted that genetic differentiation between maternal and paternal relatives is a general problem with pedigree reconstruction (Huisman, 2017; Hillary et al., 2018). Therefore, incorporating the uncertainty of differentiation or modifying the theory with the use of HS (not MHS) remains a task for future research.

As a first step in developing unbiased estimators of *N*_e_, we examined a relatively simple situation and ignored the complex but important features required in practical situations, including errors associated with kinship detection (as noted in the previous paragraph) and non-random sampling. It is expected that if non-random sampling is caused by a family-correlated sampling scheme, the effective mother size is underestimated because MHS pairs are more likely to be sampled with this sampling scheme than with random sampling. To reduce this bias, the sampling time and location should be varied, or sampling at an early life stage after hatching should be avoided; this may reduce the effect of family-correlated movement that is not addressed in the current theoretical framework.

Contemporary effective population size can provide not only an understanding of genetic health but also an indication of adult size. If the effect of overdispersion *c* is invariant across years, 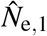 may behave as an index of the number of mothers per year, making it possible to determine the temporal trends, since Equation 11 can be rewritten as

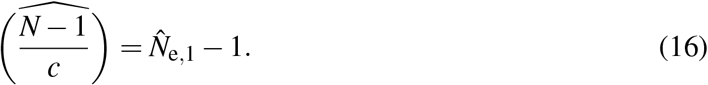

In this case, the proposed index becomes highly informative, particularly for integrating stock assessment in fisheries management using many types of data (e.g., catch data and abundance index data); this leads to more accurate estimation due to the use of fishery-independent data (Ovenden et al., 2015). Recently, Akita (2018) developed a summary statistic that indicates the degree of overdispersion; this statistic is based on the number of MHS pairs and mother–offspring pairs in the sample. The temporal trend of this statistic provides information on whether *c* is invariant across the years and thus provides criteria for determining whether 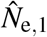 behaves as an index of the number of mothers in a population.

Finally, we note the theoretical connection of our results to the ratio of effective mother size to census size, *N*_e_/*N*. A number of studies have demonstrated that the ratio of the effective size to the census size (including fathers) in high-fecundity marine species is estimated to fall within 10^−3^−10^−6^ (Hauser and Carvalho, 2008). In our derivation, *N*_e_/*N* is approximately equal to 1/*c*. If there is no non-parental variation (i.e., 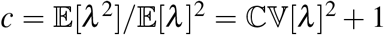), *c* cannot have a large value; thus, the ratio cannot become very small (e.g., < 10^−3^). This theoretical consideration suggests a dominant contribution of non-parental variation to a very small *N*_e_/*N*, which is consistent with the result in Waples (2016).

## Appendix

### A Probability density function and its moment of *λ*

As noted in the main text, our modeling framework does not require the specific form of *f* (*λ*); instead, it only requires the ratio of the second moment to the squared first moment 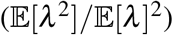. However, the specific form is required for illustrative purposes (Fig. 2a) and for the evaluation of the theoretical results (i.e., calculating the moment or running the IBM). Here, we model an age-structured fish population, which serves as a representative example, demonstrating both parental and non-parental variations.

Suppose that the mean fecundity of a mother depends on her age. Let *λ_a_* be mean fecundity, which is a function of age (denoted by *a*). The moment can be described as 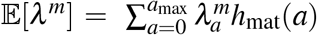, where *h*_mat_(*a*) is the frequency of mature mothers at a given age, and *a*_max_ is the maximum age. Thus, we can numerically obtain the moment from *λ_a_* and *h*_mat_(*a*).

For marine species with a type-III survivorship curve, it is generally assumed that individual fecundity is proportional to weight. Using the von Bertalanffy growth equation for body weight, *λ_a_* is explicitly described as a function of age as follows:

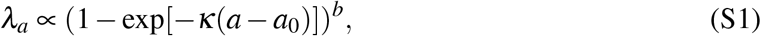

where *κ*, *a*_0_, and *b* are conventionally used parameters in the von Bertalanffy equation and represent the growth rate, the adjuster of the equation for the initial size of the animal, and the allometric growth parameter, respectively. For obtaining a specific value of *λ*, a coefficient value of 10 multiplied by the right-hand side of Eq. S1 was used when running the IBM.

The frequency of mature mothers at a given age can be written as follows:

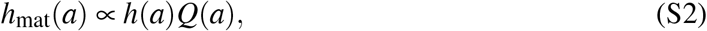

satisfying 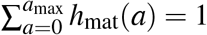, where *h*(*a*) and *Q*(*a*) represent the frequency and maturity at a given age, respectively. Although *f* (*a*) is affected by historical population dynamics and age-dependent survival, for simplicity, the mortality rate is assumed to be constant (i.e., age independent):

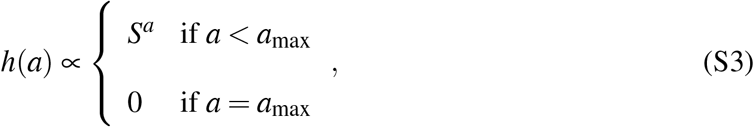

where *S* is survival probability. Maturity at age (*Q*(*a*)) is assumed to be a knife-edge function, given by

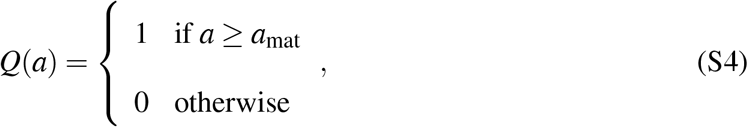

where *a*_mat_ is the mature age.

For calculating 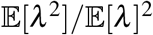, the required parameter set is (*a*_max_, *κ, a*_0_, *b, S, a*_mat_). In this paper, for the purpose of representation, we fixed the value of several parameters as follows: *a*_max_ = 20, *κ* = 0.3, *a*_0_ = 0, *S* = 0.5 and *a*_mat_ = 0. In addition, we selected parameter value 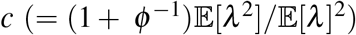 to be 1 and 10 for comparison with the results in the main text, which are derived from the parameter set (*ϕ,b*) = (1000,0.0009) and (0.1302,0.9), respectively.

Finally, we provide the specific forms of *f* (*λ*) and Pr[*k*], which are presented in the main text (Fig. 2). When *λ_a_* and *h*_mat_(*a*) are obtained, *f* (*λ*) is given by

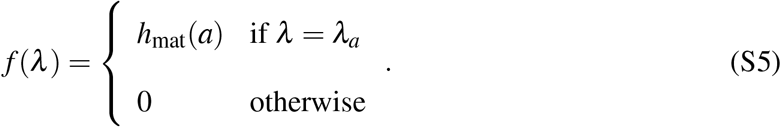

Using Eqs. 1 and S5, we can obtain the specific form of the marginal distribution of *k* by

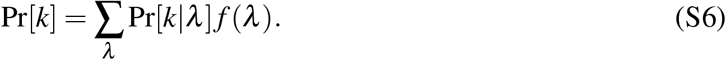

### B Probability that two offspring share an MHS relationship

Given the realized number of offspring *k*_1_,*k*_2_,…, *k_N_*, the probability that two randomly selected offspring are born to mother *i* is 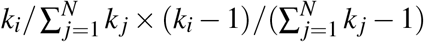. Thus, the conditional probability that two offspring share an MHS relationship with an arbitrary mother is

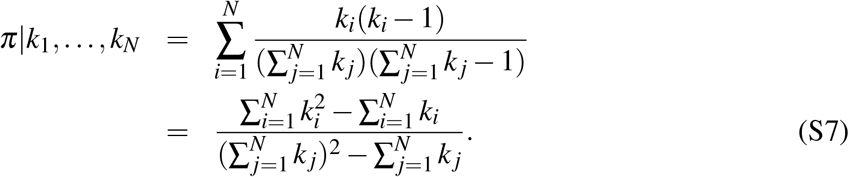

It should be noted that *k_i_* is a random variable followed by a negative binomial distribution (Eq. 1), in which the parameter of the distribution, *λ_i_*, is also a random variable followed by an arbitral function *f* (*λ*). By taking the expectation over the distribution of offspring number, the conditional probability is given by

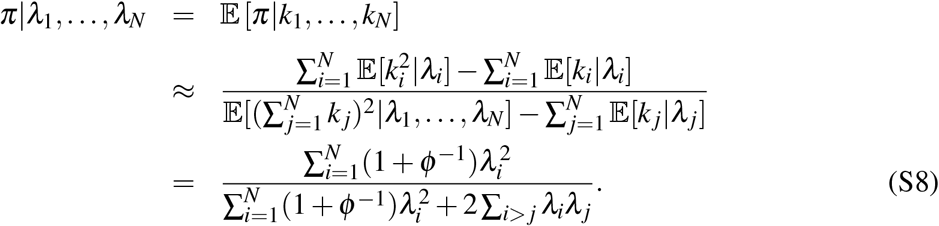

From the first to the second line, we use the approximation that 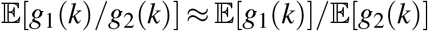. The expectations are averaged over *k* or *k*^2^, conditional on *λ*. By taking the expectation over *λ* and applying a similar approximation, the unconditional probability is given by

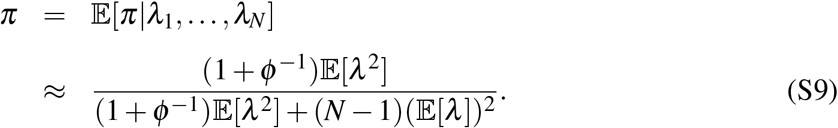

This provides the formulation described in Eq. 3. In computing the expectation, we remove the subscript (*i* or *j*) because *λ* is independent and identically distributed.

### C Properties of moment estimator of *N*

Here, we demonstrate that 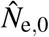 in Eq. 9 is the maximum likelihood estimator and that 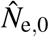 is upwardly biased, especially when *n* is small. Let *L* be the likelihood of the distribution of *H* (Eq. 6). Given the observation (i.e., *H*_obs_), the partial derivative of the log-likelihood with respect to *π* is given by

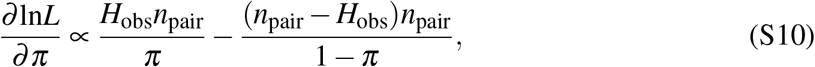

leading to the maximum likelihood estimator of *π*:

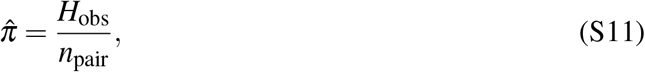

where 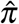 satisfies 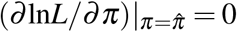. Substituting Eq. 3 into Eq. S11, we can obtain the estimator 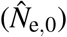 described in Eq. 9.

Consider the bias of 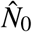 defined by 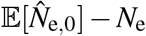. We set the following equations:

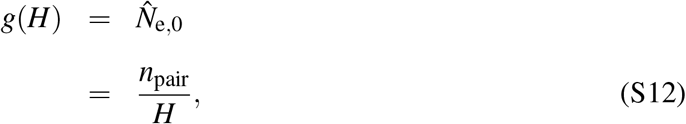

and

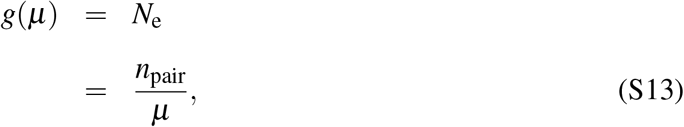

where 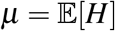. Using Eqs. 7 and 8, the bias is given by

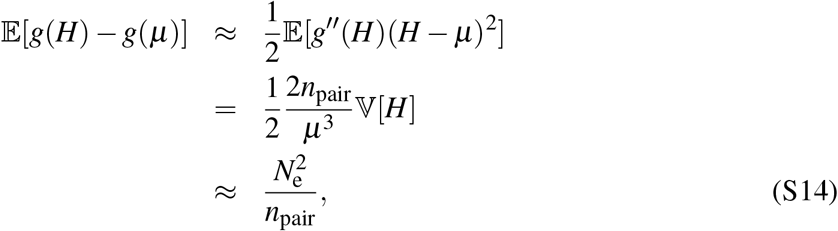

where a quadratic approximation for *g* centered at *μ* and 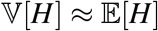 is used. The value of the right-hand side of Eq. S14 is illustrated in Figs. 3a and 3b as the bias of 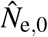.

### D Derivation of nearly unbiased estimator of *N*_e_

We consider the following

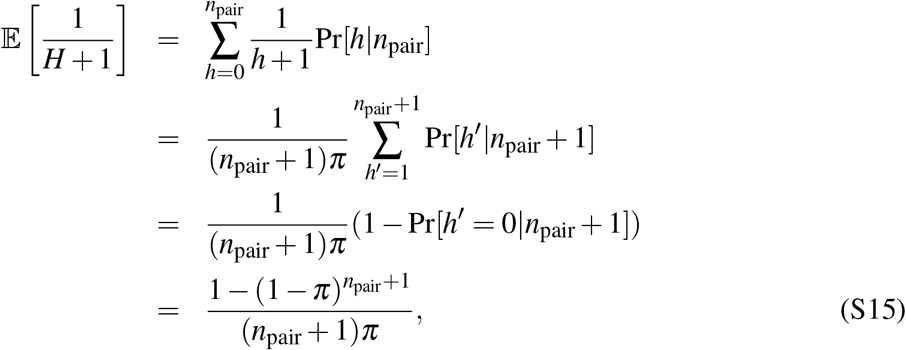

assuming the binomial form of *H* (Eq. 6). Equation S15 is not directly applied for the derivation of the estimator of *N*_e_ due to the complex formulation. Thus, we simplify the formulation as follows:

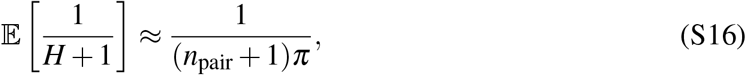

assuming that

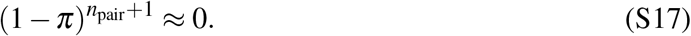

This simplification deviates from the prediction by Eq. S15 when *n* is relatively small. Replacing 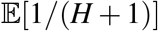 by 1/(*H*_obs_ + 1) in the left-hand side of Eq. S16, we can obtain the estimator 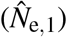 described in Eq. 11.

For the evaluation of 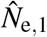, the bias is calculated. 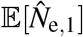 is required for the calculation and given by

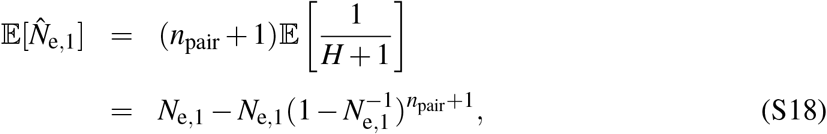

where the relationship in Eq. S15 is used. This provides the formulation of the bias, as described in Eq. 12.

### E Derivation of estimator of variance of 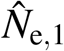

Let *ν* be the estimator of the variance of 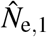. It is desirable for *ν* to be defined such that the bias (denoted *b*_var_) is reasonably small. From Eq. 11, the variance of 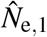 is given by

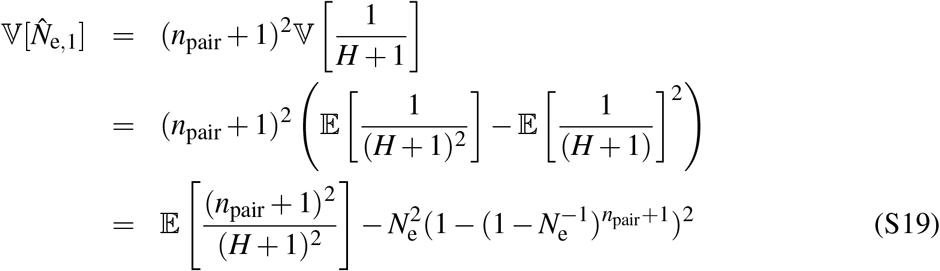

where the term 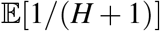 is calculated from the relationship in Eq. S15. Roughly speaking, 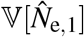 is dominated by two terms when *n*_pair_ is relatively large: 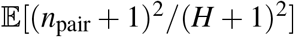 and 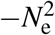. Thus, it is expected that 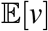 includes both terms for a reasonably small bias. We propose the following formulation for *ν*:

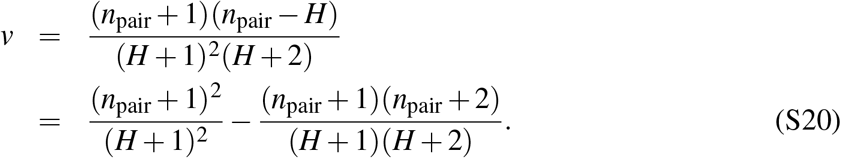

The expectation of the second term in Eq. S20 is given by

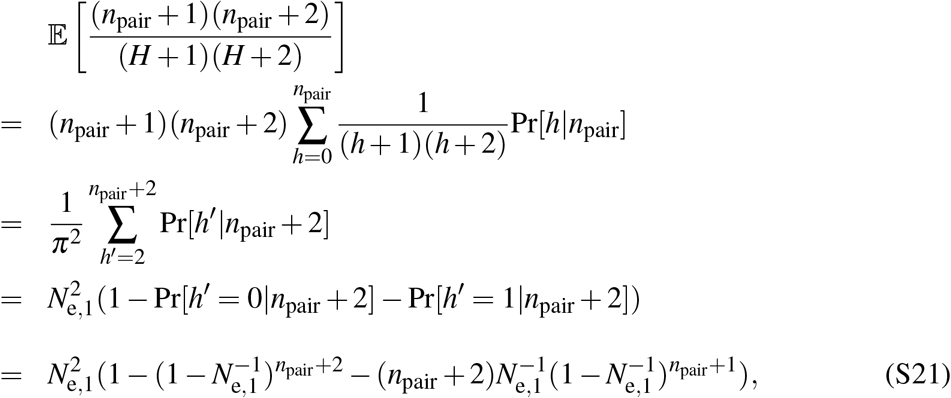

leading to a relatively small *b*_var_ when *n*_pair_ is large, which is described in Eq. 14.

## Supplementary Information

**Figure S1:**
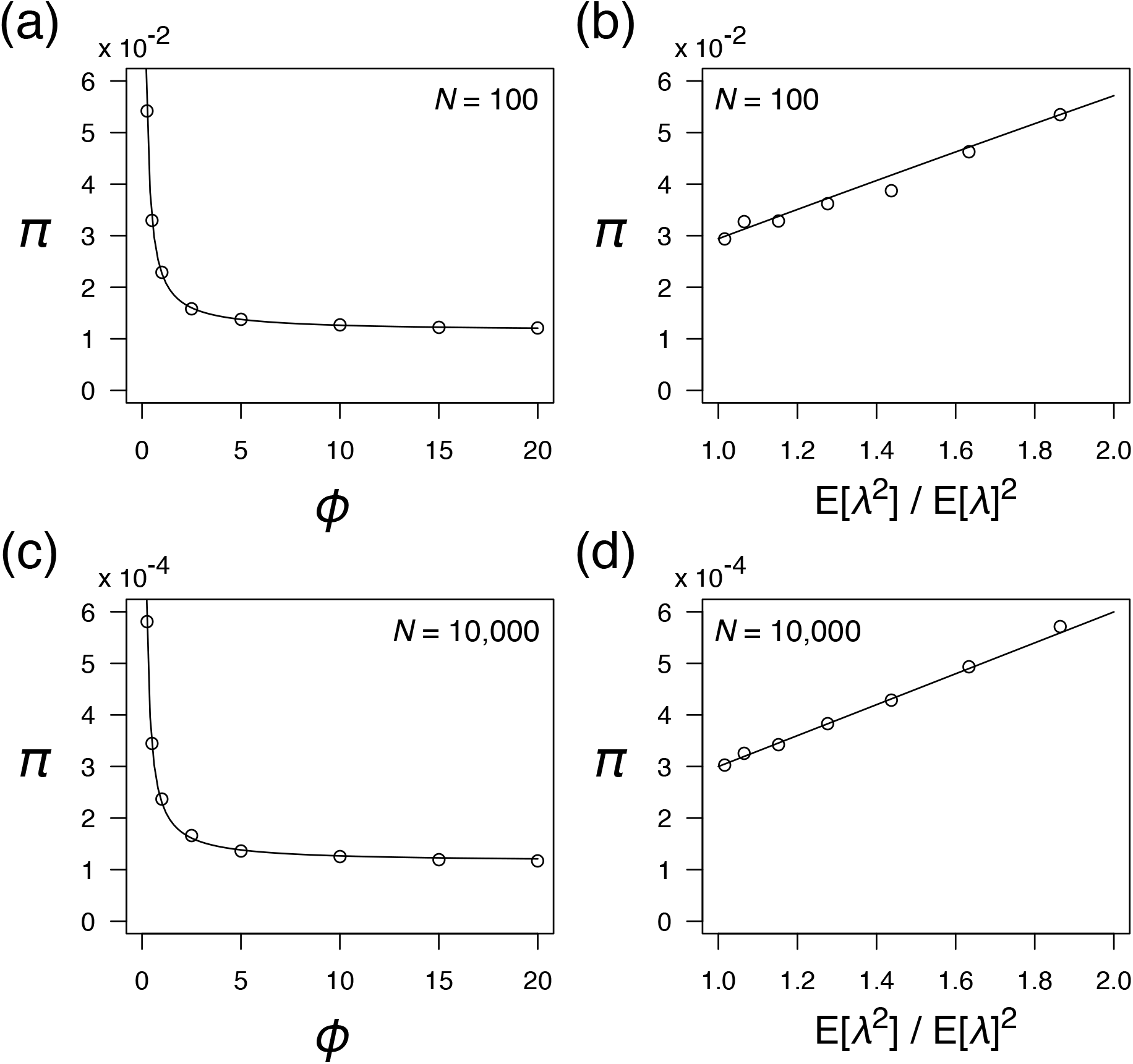
Accuracy of approximation for *π* as a function of (a), (c) *ϕ* and (b), (d) 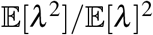. The thin line represents the approximated values (Eq. 3), and the points represent the simulated values from 10,000,000 replications. (b), (d) 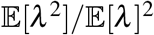 is calculated from the distribution of *λ* (see Appendix A) with *b* = 0.3,0.6,0.9,1.2,1.5,1.8, and 2.1. (a) *b* = 0.9. (b) *ϕ* = 0.5. The definition of parameter *b* is described in Appendix A.

**Figure S2:**
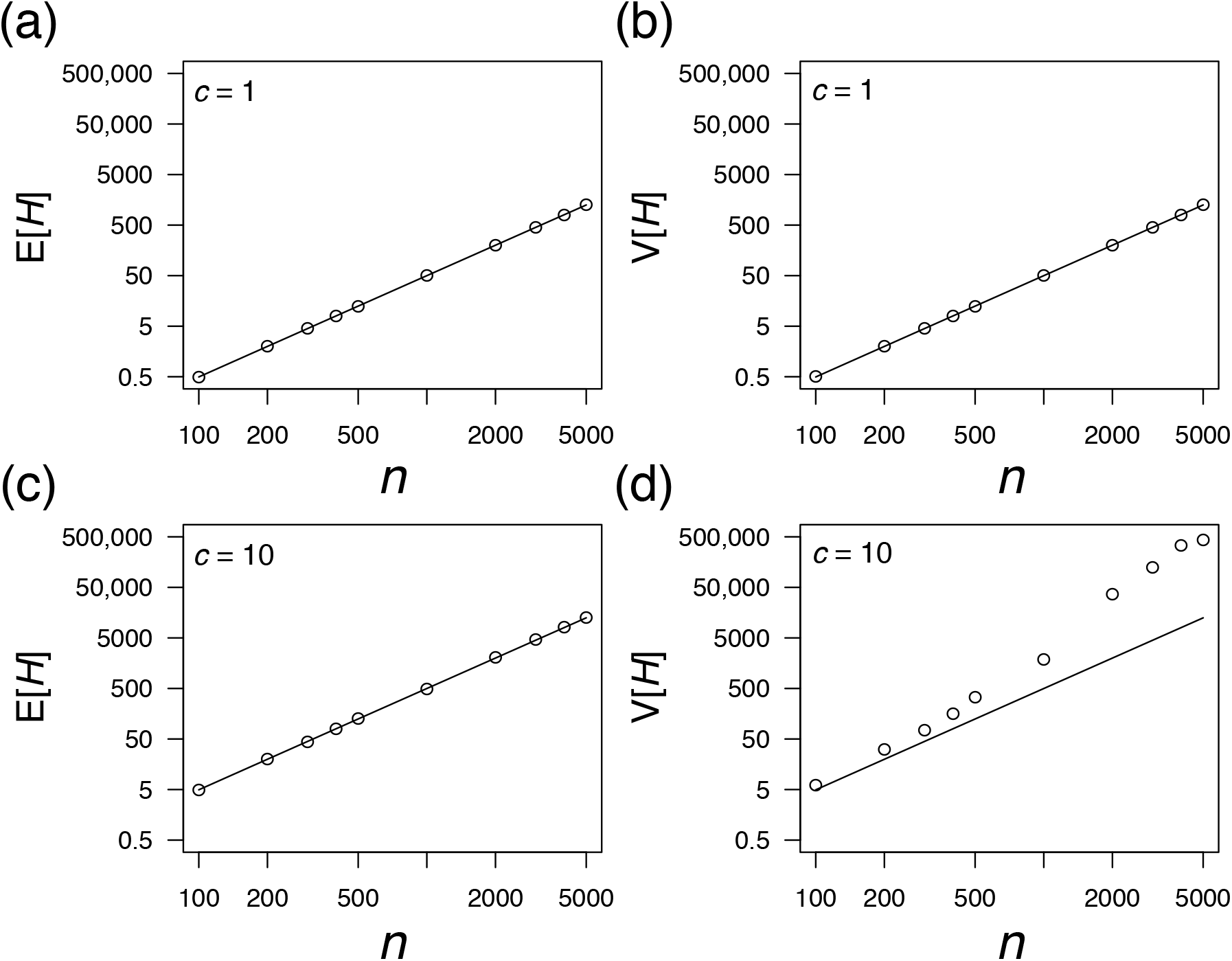
Accuracy of theoretical prediction for (a), (b) 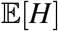 and (c), (d) 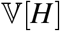 as a function of *n*. Thin lines indicate theoretical values (see Eq.s 7 and 8) and points are obtained from simulated data (1,000,000 replications). The value of *c* is indicated in the legend. *N* = 10,000. Both the *x*-axis and *y*-axis are log-scale.

**Figure S3:**
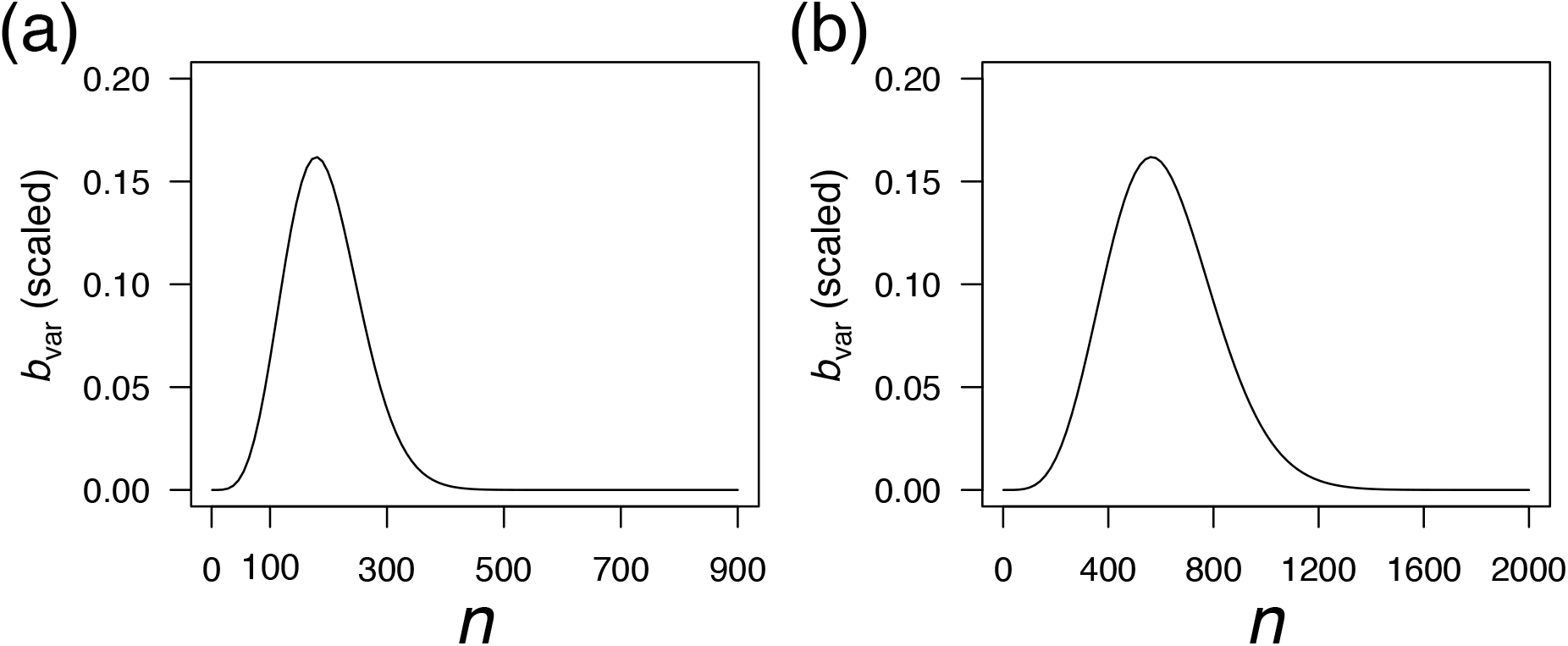
Bias of 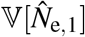 as a function of *n*. The value of the bias is scaled by 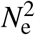. (a) *N*_e_ = 10,000. (b) *N*_e_ = 100,000.

